# Field-validation of multiple species distribution models shows variation in performance for predicting *Aedes albopictus* distributions at the invasion edge

**DOI:** 10.1101/2025.05.01.651656

**Authors:** Anna V. Shattuck, Brandon D. Hollingsworth, Jared Skrotzki, Scott R. Campbell, Christopher L. Romano, Courtney C. Murdock

## Abstract

Climate and land use changes have resulted in range expansion of many species. In this shifting disease landscape, it is important to leverage tools that can predict the potential distributions of invading vectors to target surveillance and control efforts and identify at risk populations. Species Distribution Models (SDMs) are widely used to predict ranges of invasive species; however, invasive species often violate assumptions of equilibrium and niche conservatism. Moreover, these studies are rarely validated using independent data. Here, we use long-term mosquito surveillance data for *Aedes albopictus*, a highly invasive mosquito capable of transmitting several arboviruses, at its range-edge to evaluate a variety of SDMs (MaxEnt, GAM, Random Forest, Boosted Regression Tree) in predicting *Ae. albopictus* range. We identify key environmental drivers of distributions and areas where models tended to disagree in predicting occurrence. At sites where models disagree, we sampled for *Ae. albopictus* to generate an independent dataset for field-validation of models in addition to the common practice of cross-validation. Finally, we determine if models based on early invasion data can predict later stage invasion ranges. We found that landscape and climatic variables are important drivers of population distributions. SDM methods varied in predictive accuracy between models and across validation methods (i.e. cross-vs. field-validation). GAM and MaxEnt best predicted later-stage invasion distributions, requiring fewer years of training data. Our work shows that SDMs can be useful tools to predict the ranges of invasive species and highlights the importance of comparing predictions of invasive species’ range.

**Author Summary:** Mosquitoes are greatly impacted by their surrounding environment. Environmental changes such as those driven by climate change or changes in land use (i.e. urbanization, etc.) can have profound impacts on mosquito ranges. Species distribution models (SDMs), which use the occurrence data of a species in combination with environmental data (e.g., landscape characteristics, temperature, etc.) to estimate suitable habitats for the species, are helpful tools to understand how species ranges can change with the environment through time and space. There are many different species distribution models to choose from – each with different methods for estimating suitable habitat – making it important to compare the performance of different models. Furthermore, because invasive species often violate assumptions of SDMs, it is important to rigorously explore how different methods perform when predicting invasive species ranges in newly invaded areas. In this study, we tested the predictive accuracy of different species distribution models, and we found it varied across modeling and validation method. We also tested whether models could use early-stage invasion data to predict the late-stage invasion distributions of *Ae. albopictus.* We found that some modeling methods performed well while others needed more data to improve accuracy of late-stage invasion range predictions. In summary, when modeling species ranges using SDMs, it is important to use multiple methods and compare results because methods will likely disagree, and further sampling may be required to determine which model is most accurate.

## Introduction

Mosquito-borne disease is a significant public health threat and economic burden both world-wide and within the United States [1, 2]. In recent decades, there has been an increase in mosquito-borne disease incidence, despite widespread control efforts [3–6]. The complex and often combined effects of climate and land-use change have resulted in the emergence and re-emergence of pathogens (chikungunya, Zika, malaria, yellow fever, Oropouche) and the expansion of many vector ranges outside native distributions [7–11]. Whether distributions of mosquito species shift with climate or land use change will likely be due to ecological and physiological factors [12, 13]. Recent expansions of invasive mosquitoes include *Aedes albopictus*, an ecologically flexible vector of several arboviruses, whose range is continuing to expand northward in response to climate warming [12, 14], and *Anopheles stephensi*, the urban malaria vector of South Asia, into the horn of Africa [15, 16]. In this shifting disease landscape, it is increasingly important to accurately predict the potential distributions of invading vectors and identify at risk human populations.

Species Distribution Models (SDMs), also referred to as Ecological Niche Models, are widely used to characterize the ecological niches and distributions of species from a variety of taxa [17–21]. These models leverage species occurrence data in combination with environmental data (e.g., landscape characteristics, urbanization, abiotic variables) to estimate suitable habitats for species of interest [19, 22, 23]. Several SDMs are available, including machine learning (e.g., MaxEnt, Boosted Regression Tree, Random Forest), statistical (e.g., Generalized Additive Models), and mechanistic approaches (e.g., thermal performance) [19, 23]. It is now standard practice to use multiple SDM methods or ensemble models – models that combine multiple SDM methods – to predict species distributions and compare results of different methods [18, 24, 25].

Despite the popularity of SDMs, there has been less effort to compare the predictive accuracy of different modeling methods and to assess predictive performance through field-validation. Field validation can be economically costly or logistically challenging in some scenarios (e.g., with future climate change); however, when possible, it can be used to improve an SDM’s performance and more rigorously compare and evaluate model predictions [26–29]. Cross-validation, which leaves out a portion of the dataset from training to assess a model’s predictive accuracy, provides a method for testing out-of-sample (i.e. testing model’s performance on data not included in training) or out-of-range (i.e. testing model’s performance on environmental data outside the range of environmental data used in training) predictions. However, field-validating predictions using independent datasets is critical. Using SDMs to project the final distributions of invasive species is also potentially problematic, as invasive species often violate the assumptions of equilibrium and niche conservatism in SDMs [21]. Few studies have focused on understanding the extent these models can be used to predict final distributions using early invasion data, but they have shown mixed results in the ability of SDMs to predict expansions of invasive species into novel regions [30–32].

We use 16 years (2008 to 2023) of mosquito surveillance data from Suffolk County, NY, to evaluate a variety of SDMs in predicting *Ae. albopictus* distributions across a heterogenous, and seasonally shifting, landscape. New York represents the northern edge of the current range of *Ae. albopictus* in the United States, with *Ae. albopictus* first detected in Suffolk County in 2008. Thus, this is an ideal system to test the ability of SDM approaches to predict late-stage distributions of invading species, as well as the number of years of data required to achieve high accuracy. We employed common machine learning (MaxEnt, random forests, and boosted regression trees) and statistical (generalized additive models) SDMs. Due to the limitations of statistical and machine learning approaches in predicting novel scenarios, we also employed a mechanistic temperature-dependent population growth model. We had the following objectives with this study: 1) identify the effects of key environmental drivers on distributions of *Ae. albopictus*, 2) determine model prediction accuracy based on out-of-sample cross validation and field validation, and 3) determine how early in an invasion it becomes possible to accurately predict final distributions for *Ae. albopictus*.

## Methods

### Study area

Suffolk County, NY, is the easternmost county of Long Island located less than 100 km east of New York City. It contains a heterogeneous landscape, including highly urbanized, agricultural, forested, and sandy areas. *Ae. albopictus* was first detected in Suffolk County in 2004 but was not detected again until 2008, when it was likely established, having since spread throughout the county [33]. Currently, southeastern New York and Connecticut constitute the northernmost range edge of *Ae. albopictus* in the United States. However, predictions suggest that with climate change (warming temperatures, land-use change), *Ae. albopictus* will continue to expand its range [12, 14, 34–36].

### Data

We used long-term adult mosquito surveillance data collected by the Suffolk County Arthropod-Borne Disease Laboratory from 2008 to 2023 to train SDMs. These sites were sampled weekly using CDC light (John W. Hock Company, Gainesville, Florida, USA) and CDC gravid traps (John W. Hock Company, Gainesville, Florida, USA) at established sites throughout the county from May to October, with additional sites trapped in response to reported vector-borne disease cases and West Nile virus positive dead birds. All collected adult mosquitoes were identified to the lowest taxonomic level possible.

All SDMs had *Ae. albopictus* presence/absence as the response variable, and we chose five environmental variables as predictors based on the ecology of *Ae. albopictus* [37–40]. Day and night land surface temperatures aggregated at a 1km^2^ scale [41], enhanced vegetative index (EVI) aggregated at a 0.25km^2^ scale [42], percent impervious surface (fractional impervious surface) aggregated at a 0.25km^2^ scale [43], and land cover classifications at a resolution of 0.25km^2^ [43]. Missing data was predicted using linear interpolation between the two closest data points spatially or temporally. Environmental information from 2023 was used to predict distributions of *Ae. albopictus* in 2024. We assumed changes in land characteristic data (i.e. percent impervious surface, land cover classification) would be minimal between years, and the variables that had more seasonal fluctuation in value (i.e. temperature, EVI) would be similar between years with minimal effects of long-term trends such as climate warming. To account for the ongoing invasion, we used convex hulls with a buffer of 4000 m from each site point to restrict the geographic extent each year.

### Training SDMs

We applied four widely used SDMs to our surveillance dataset: Generalized Additive Models (GAMs), Maximum Entropy (MaxEnt), Random Forest (RF), and Boosted Regression Tree (BRT) [44–47]. GAMs apply flexible, non-linear relationships between the response variable (species occurrence) and explanatory variables (environmental covariates) to capture distribution patterns. MaxEnt, a presence-only model, uses a maximum entropy approach to estimate the probability of a phenomenon occurring (i.e. a species being present). RF and BRT are both ensemble methods using decision trees. RF builds decision trees independently, each using a subset of the dataset, then combines each individual prediction to create a final prediction. BRT, in contrast, builds decision trees sequentially, using a boosting algorithm to minimize losses in predictive performance. We used the same predictors across modeling methods and trained with the same dataset of *Ae. albopictus* surveillance data from 2008-2023. GAMs were fit with k = 5 to avoid overfitting. All models were fit in R [48] using the *randomForest* package [47] for RF; *sdm* package [45] for MaxEnt and BRT; and *mgcv* package for GAM [46]. We also used a mechanistic SDM (*λ*) based on temperature (T)-dependent population growth (Equation 1).

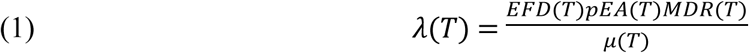

This mechanistic model includes information relevant to mosquito population growth and decline including eggs per female per day (EFD), mosquito development rate (MDR), probability of egg to adult survival (pEA), and adult mortality rate (*μ*). We used the thermal performance curves and parameter estimates for these temperature dependent life history traits from [49].

Model accuracy was first estimated by calculating the area under the receiver-operator curve (AUC) based on out-of-sample cross-validation using the long-term surveillance dataset. We evaluated both out-of-sample and out-of-range (i.e. models ability to predict beyond the range of environmental variables used in training) predictive ability by sub-setting the dataset 50 times into training (80%) and testing (20%) groups (out-of-sample) and withholding each township (n=10) in succession (out-of-range). Random sampling was employed to assess the ability of models to interpolate spatially and temporally, whereas township sampling was employed to assess the ability of models to extrapolate into geographically (and potentially environmentally) novel regions.

### Selecting Field Sites Based on Model Disagreement

We selected field sites based on SDM disagreement in predictions of habitat suitability (Fig. 1). The mechanistic model was not included in calculations of model disagreement and field-validation as cross-validation showed overall poor predictive performance (AUC=0.538). Predictions of habitat suitability were made for each month of sampling (July-September) in 2024, representing early, peak, and late season for *Ae. albopictus* activity, respectively. Models were trained on the full surveillance dataset. Then, using out-of-sample ROC curves, we choose suitability thresholds for each model that maintained the sensitivity (true presence) and specificity (true absence) for *Ae. albopictus*. Since models predict a continuous variable, threshold represents the cutoff line that dictates whether the prediction for a site is positive or negative for *Ae. albopictus*.

**Figure 1.**
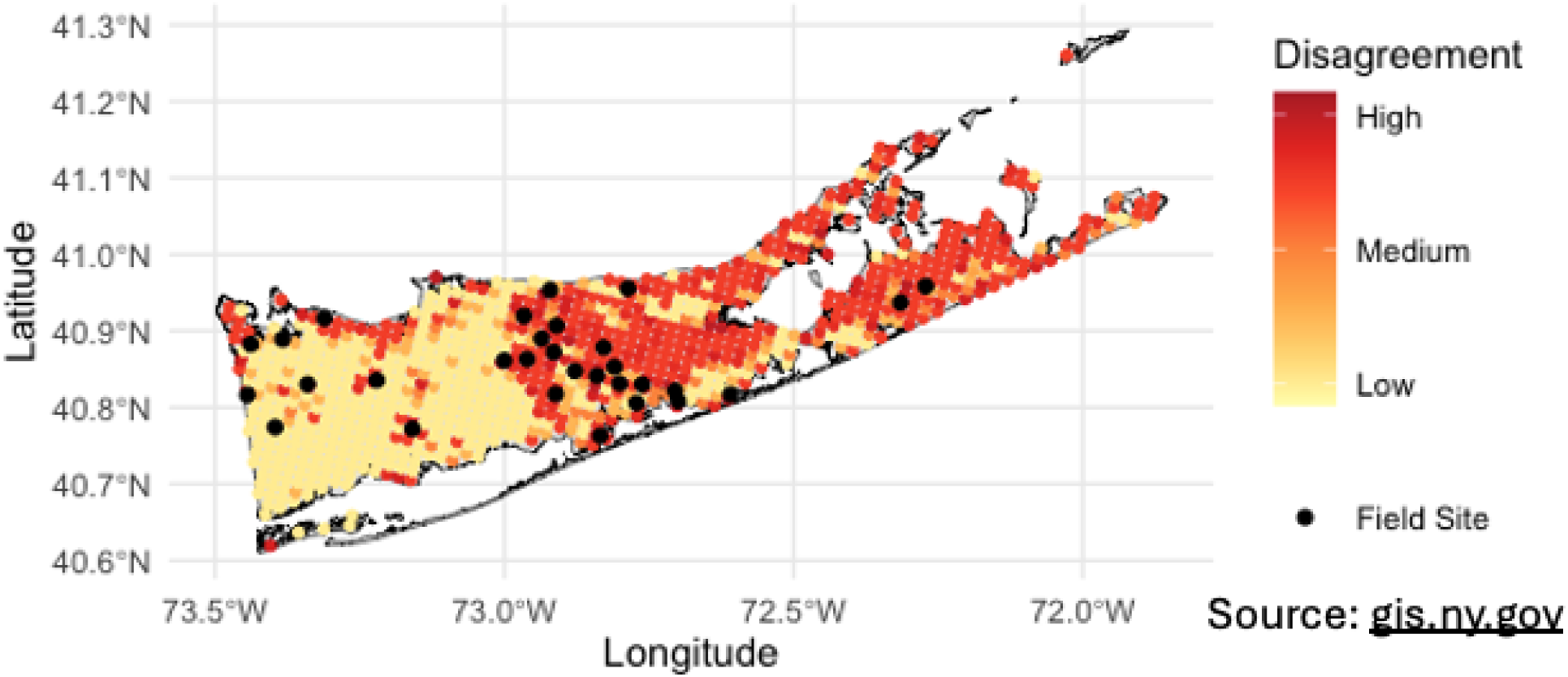
Map of disagreement between SDMs and of field sites in Suffolk County, NY, USA. Map of Suffolk County, New York, U.S.A. with the degree of disagreement among predictions for the presence of *Aedes albopictus* displayed. Areas of high disagreement among models are colored red with areas of low disagreement colored as yellow. Black points represent the field sites (n = 30) that were chosen based on high disagreement among models. Suffolk County shape file from NYS GIS Civil Boundaries Program (https://gis.ny.gov/civil-boundaries).

Model disagreement was calculated as the total number of disagreements arising from all pairwise combinations of model predictions (Equation 2, Fig. 1).

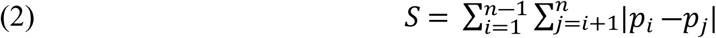

Let *i* and *j* be indices representing different models, where *i*, *j* ∈ {1, 2, …, *n*} and *i* < *j*, *p* is the prediction of each model, and *S* the total disagreement across models. Field sites (n = 30) were selected based on total disagreement across July-September and our ability to access the site and safely set up a mosquito trap (Fig. 1). Sites had not been sampled previously by the Suffolk County Arthropod-Borne Disease Laboratory.

### Field Sampling

We sampled *Ae. albopictus* using BG-Sentinel traps (Biogents AG, Regensburg, Germany) baited with BG-Lure Cartridges [50]. At each site, one trap was set in total shade for 48 hours, catch bags were collected and replaced every 24 hours, and all traps were set once monthly from July-September 2024. Trap nights were removed from the dataset if it was clear the trap had been tampered with (e.g., wildlife encounter). This resulted in 177 total sampling events out of 180 possible sampling events, with one sampling event removed in July and two in September. Mosquitoes were identified to species level under dissection microscopes at the Suffolk County Arthropod-borne Disease Laboratory using morphological keys [51, 52].

### Exploring the Effect of Additional Training Data on SDM Predictive Abilities

To determine how the addition of more years of surveillance data from early in an invasion improved the predictive accuracy of models attempting to forecast *Ae. albopictus* distributions (S1 Figure), we applied a modified expanding window training approach to the models described above (MaxEnt, GAMs, RF, BRT) where the testing dataset remained constant (disagreement sites in July-September 2024). The training process for each model involved incrementally incorporating an additional year of training data chronologically (e.g., iteration 1 training dataset = 2008, iteration 2 training dataset = 2008-2009, etc.; S2 Figure) until all years of data were incorporated. At the end of the training process, we produced 16 prediction sets for each site averaging across months (1 prediction set per iteration*16 iterations) of 2024 habitat suitability from each of the four models (30 sites * 16 iterations * 4 models).

### Evaluating SDM Performance in Areas of Disagreement

We estimated the predictive skill of RF, BRT, GAM, and MaxEnt models using the logarithmic score, a strictly proper scoring metric that assigns scores based on how close model predictions are to the observed outcome [53]. The logarithmic score (Equation 3) was calculated using the observed occurrence (x = {0,1}) at a field site and the predicted suitability (*p* = [0,1]). Here, a lower score represents higher predictive skill.

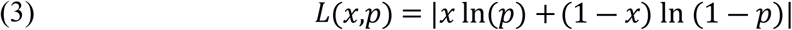

The logarithmic score was calculated for each site across each month (July-September 2024) and then averaged for each of the four SDM methods per month. We used the logarithmic score to assess the predictive accuracy of each SDM trained on the entire surveillance dataset and on the modified expanding window training datasets.

### Statistical Analysis

All statistical analysis was conducted in R version 4.3.2 using the *stats* package [48]. To explore differences in estimated suitability between modeling methods and across a mosquito season, we tested the effects of modeling method, month, and their interaction on estimated habitat suitability using ANOVA with a Tukey post-hoc test to explore pairwise differences. Habitat suitability was estimated for a 2024 prediction data frame (described in previous section) by MaxEnt, BRT, RF, and GAM that were trained on the entire surveillance dataset. As we were also interested in whether models disagreed more in specific land cover types, we tested the effect of land use classification on model disagreement using an ANOVA followed by a Tukey post-hoc test to explore pairwise differences. Finally, we tested for differences in logarithmic scores between models across months using an ANOVA followed by a Tukey post-hoc test to address pairwise differences.

To determine the effect of environmental covariates on habitat suitability, we created partial dependence plots (PDPs) for the following predictors: impervious surface, EVI, day land surface temperature, night land surface temperature, land cover, month, and year. PDPs show marginal effects of predictors on the response variable. To create a PDP for an environmental covariate, we held other environmental covariates constant at their average value or develop – low, in the case of land cover classification, and varied the covariate of interest. We then allowed each model that was trained on the full surveillance dataset to predict habitat suitability. We did this process for each environmental covariate.

## Results

### Habitat Suitability

Predicted habitat suitability for *Ae. albopictus* varied spatially and seasonally in Suffolk County, NY (Fig. 2, Fig. 3). Habitat suitability was predicted to be greatest in the western half of Suffolk County. This area is more urban and experienced warmer day and night temperatures (Fig. 2, S3 Figure). Lowest suitability was predicted in the middle portion of the island (between 73.0°W and 72.5°W), where the Central Pine Barrens is located, an area of publicly protected land that houses what remains of the Atlantic coastal pine barrens ecoregion. This was expected because the Central Pine Barrens has a low human population, sandy, well-draining soil, and is dominated by pines.

**Figure 2.**
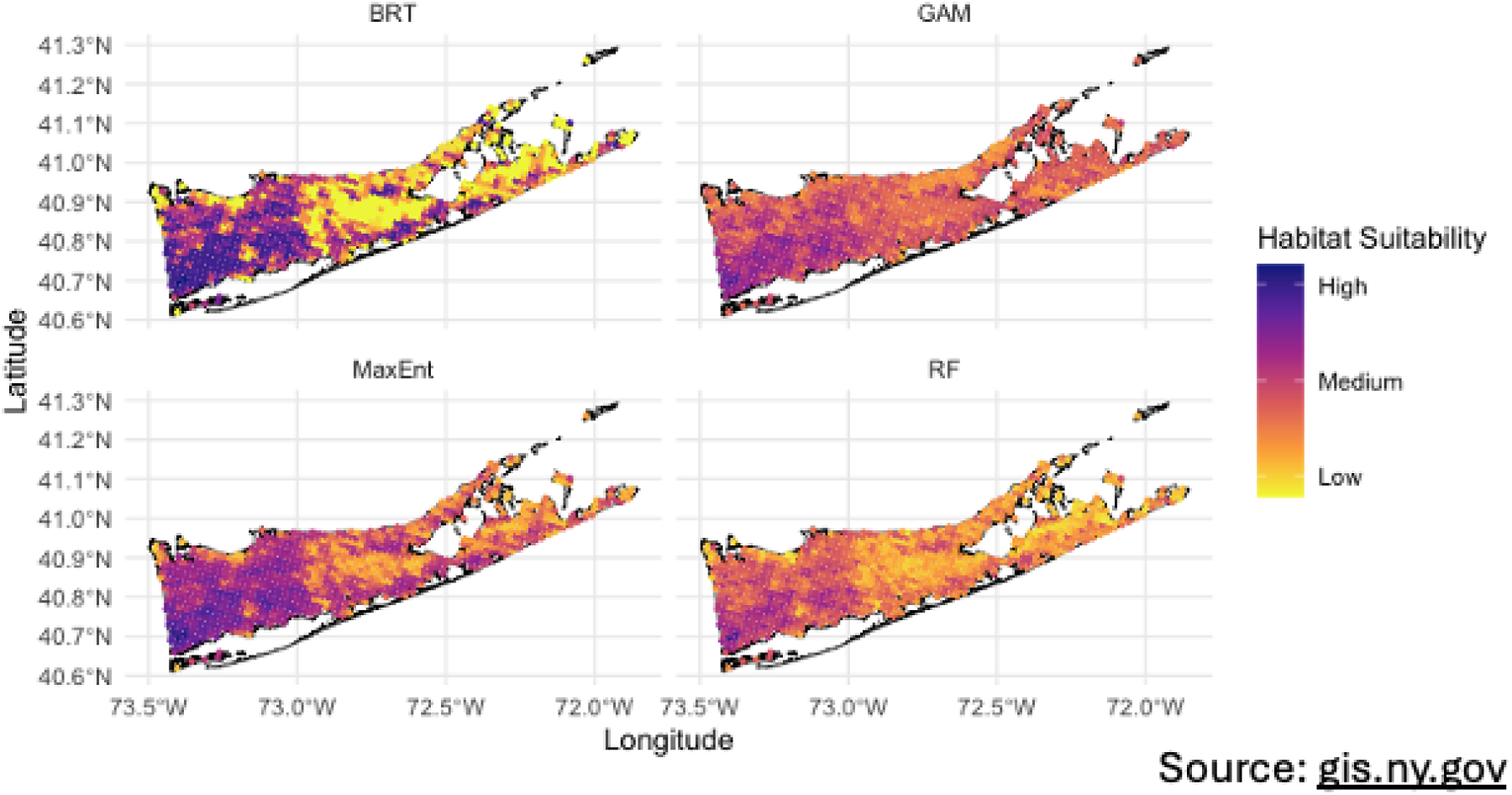
Maps of habitat suitability for *Ae. albopictus* across SDMs in Suffolk County, NY, USA. Mean predicted habitat suitability for *Aedes albopictus* across May to October for Suffolk County, New York, U.S.A. by four species distribution modeling (SDM) approaches (BRT = Boosted Regression Tree; GAM = Generalized Additive Model; MaxEnt = Maximum Entropy; and RF = Random Forest). Suitability estimates are averaged across sampling months for each SDM. Darker colors indicate higher predicted habitat suitability for *Ae albopictus*. Suffolk County shape file from NYS GIS Civil Boundaries Program (https://gis.ny.gov/civil-boundaries).

**Figure 3.**
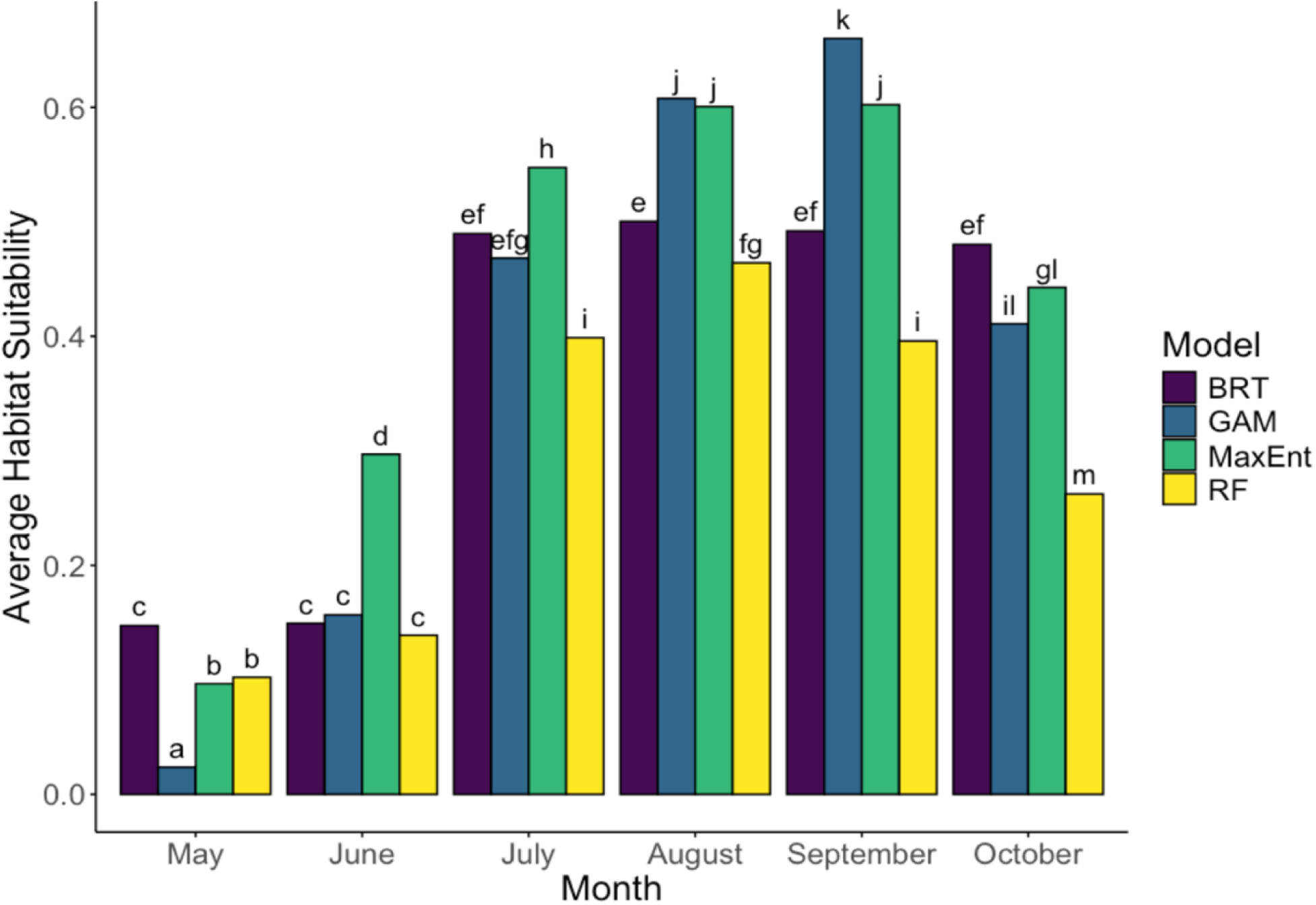
Seasonal change in estimated habitat suitability for *Ae. albopictus* across SDM methods. Average estimated habitat suitability of *Aedes albopictus* for Suffolk County, New York, U.S.A. from each species distribution model (BRT = Boosted Regression Tree, navy; GAM = Generalized Additive Model, blue; MaxEnt = Maximum Entropy, green; and RF = Random Forest, yellow). Habitat suitability is averaged across each month of the sampling season. Letters indicate significant differences (α=0.05) calculated from an ANOVA followed by a Tukey post-hoc analysis.

Greatest habitat suitability was estimated from July – September (Fig. 3). Typically, MaxEnt and GAM predicted higher mean suitability each month, while RF tended to predict the lowest mean suitability (Fig. 3). BRT predicted almost the same suitability values for May and June and then almost the same suitability values for July, August, September, and October, instead of following a more gradual increase over time as other methods did (Fig. 3).

RF (AUC = 0.778) and MaxEnt (AUC = 0.779) had the highest out-of-sample AUC, followed by BRT (AUC = 0.756), then GAM (AUC = 0.677, S4 Figure). The mechanistic model performed marginally better than what would be expected by random chance (out-of-sample AUC = 0.538, S5 Figure). The discriminatory ability of our models worsened with out-of-region predictions. GAM (AUC = 0.710), RF (AUC = 0.707), and MaxEnt (AUC = 0.710) all performed similarly, with Boosted Regression Tree performing the worst (AUC = 0.660, S6 Figure). While out-of-region values are worse than out-of-sample, AUC values in the range we are seeing are typically considered fair in terms of discriminatory ability.

### Environmental Drivers of Habitat Suitability

Modeling methods assigned different relationships between environmental predictors and suitability estimates (S7 Figure). All models assigned a positive effect of impervious surface on habitat suitability (S7 Figure). BRT, RF, and MaxEnt predicted increased habitat suitability as impervious surface increased from 0% to 40%. After 40% imperviousness, BRT and RF predicted habitat suitability to level out with further increases in impervious surface, whereas MaxEnt predicted habitat suitability would continue to increase with impervious surface cover (S7 Figure). GAM also predicted increasing habitat suitability with increased impervious surface, albeit linearly (S7 Figure). Interestingly, for EVI, methods assigned different effects on habitat suitability (S7 Figure). RF and MaxEnt assigned negative relationships between EVI and habitat suitability (S7 Figure). BRT assigned no effect of EVI on habitat suitability, while GAM assigned a positive relationship between EVI and suitability (S7 Figure). Models assigned positive relationships between day/night land surface temperature and habitat suitability, except for BRT, which found no effect (S7 Figure). Lastly, models assigned different relationships between land class and suitability. BRT found no effect of land cover on habitat suitability (S7 Figure). MaxEnt found habitat suitability to be higher in minimally developed (developed – low), altered, and natural areas than more densely development areas (developed - medium/high). RF found altered, natural, and moderate to high development (developed medium/high) to be more suitable than minimally developed habitat (developed – low) (S7 Figure). GAM found natural and minimally developed (developed – low) to have higher suitability than more developed (developed - med/high) and altered habitats (S7 Figure).

### Field-validating models in areas of disagreement

Models disagreed most often at sites where land cover was classified as natural, followed by altered, then developed – low and developed – medium/high (S8 Figure). As a result, most of the sites we selected were classified as natural (n = 21) or altered (n = 8) with a single site classified as developed – low (n = 1). The log score of models in areas of disagreement varied across modeling methods and sampling month (Fig. 4a). MaxEnt and GAM were the most accurate in predicting the occurrence of *Ae. albopictus* throughout the entire sampling season, scoring less than 0.75 and 0.85 respectively (Fig. 4a). The predictive ability of BRT and Random Forest improved from July to September by 0.62 and 0.37 points respectively (Fig. 4a). September was the only month that all models had similar performance in predicting *Ae. albopictus* presence (Fig. 4a). BRT and Random Forest consistently underpredicted *Ae. albopictus* occurrence, while GAM consistently overpredicted (Fig. 4b). MaxEnt underpredicted in July and August and overpredicted in September (Fig. 4b).

**Figure 4.**
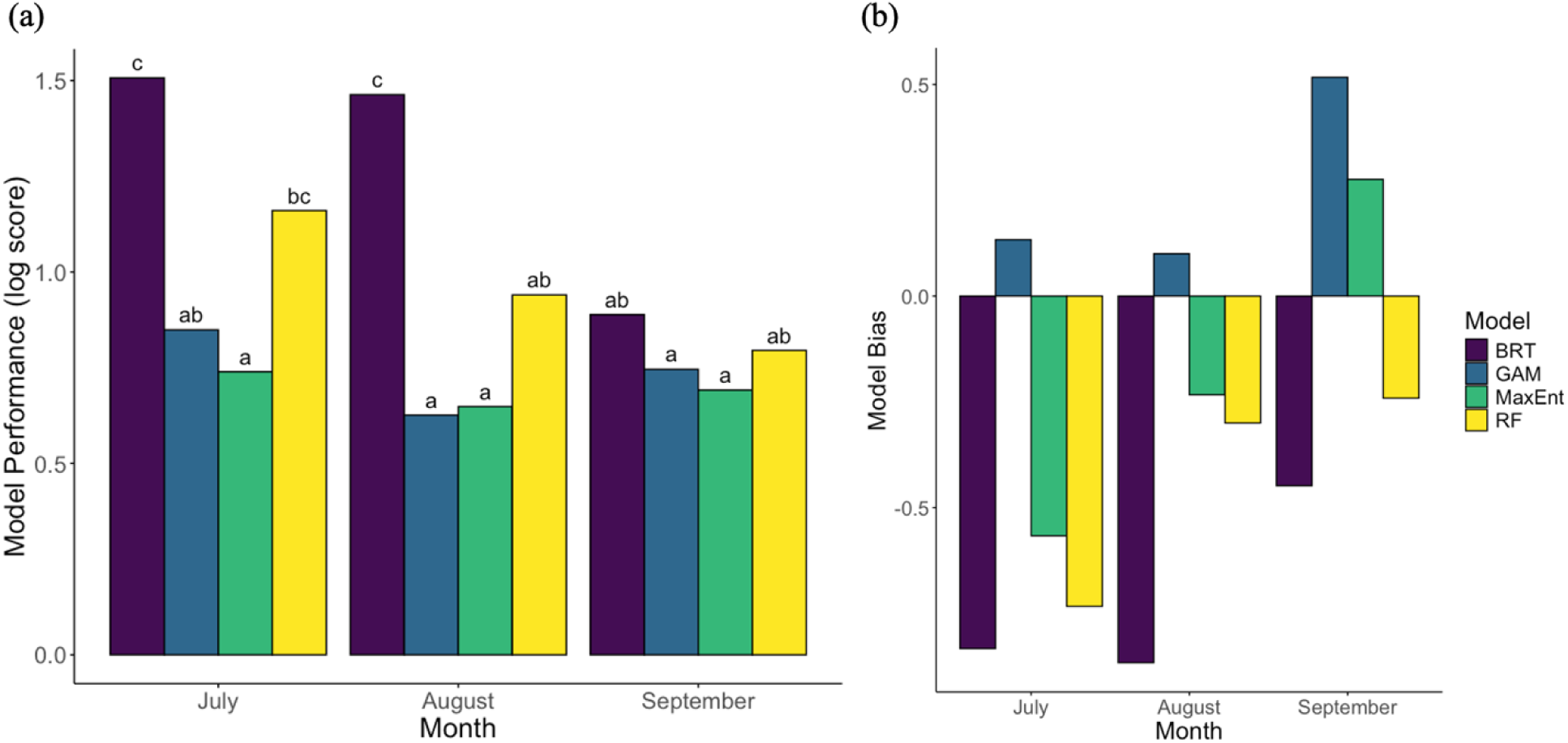
Seasonal SDM logarithmic score performance (a) and prediction biases (b) at sites of model disagreement. Graph a) represents model performance, calculated as a logarithmic score, at field sites of disagreement in predicted habitat suitability (presence) for *Ae. albopictus* in Suffolk County, New York, U.S.A. across the months of July – September. Bars represent average logarithmic scores colored by modeling method (BRT = Boosted Regression Tree, navy; GAM = Generalized Additive Model, blue; MaxEnt = Maximum Entropy, green; RF = Random Forest, yellow). Higher logarithmic scores indicate lower predictive skill. Graph b) represents the average bias of different modeling methods in predicting habitat suitability (*Aedes albopictus* presence) based on field sampling *Ae. albopictus* across 30 selected sites (Suffolk County, New York, U.S.A.) in 2024. If model bias is positive, the model is overpredicting occurrence. If model bias is negative, the model is underpredicting occurrence. Larger model biases indicate more mispredictions in a specific direction. Letters indicate significant differences (α=0.05) calculated from an ANOVA followed by a Tukey post-hoc analysis.

### Use of early invasion data in predicting establishment

Our results show that the addition of training data after 2012 did not greatly improve the overall predictive ability of our models (Fig. 5). While Boosted Regression Trees and Random Forest experience a decline in performance with the inclusion of data from 2015, the predictive ability of both models improves with additional training data from 2017 (Fig. 5). The predictive ability of GAM did not require training data past early invasion (2009) to improve predictions of species occurrence in 2024 (Fig. 5), while MaxEnt gradually improved in performance with the addition of more data each year (Fig. 5).

**Figure 5.**
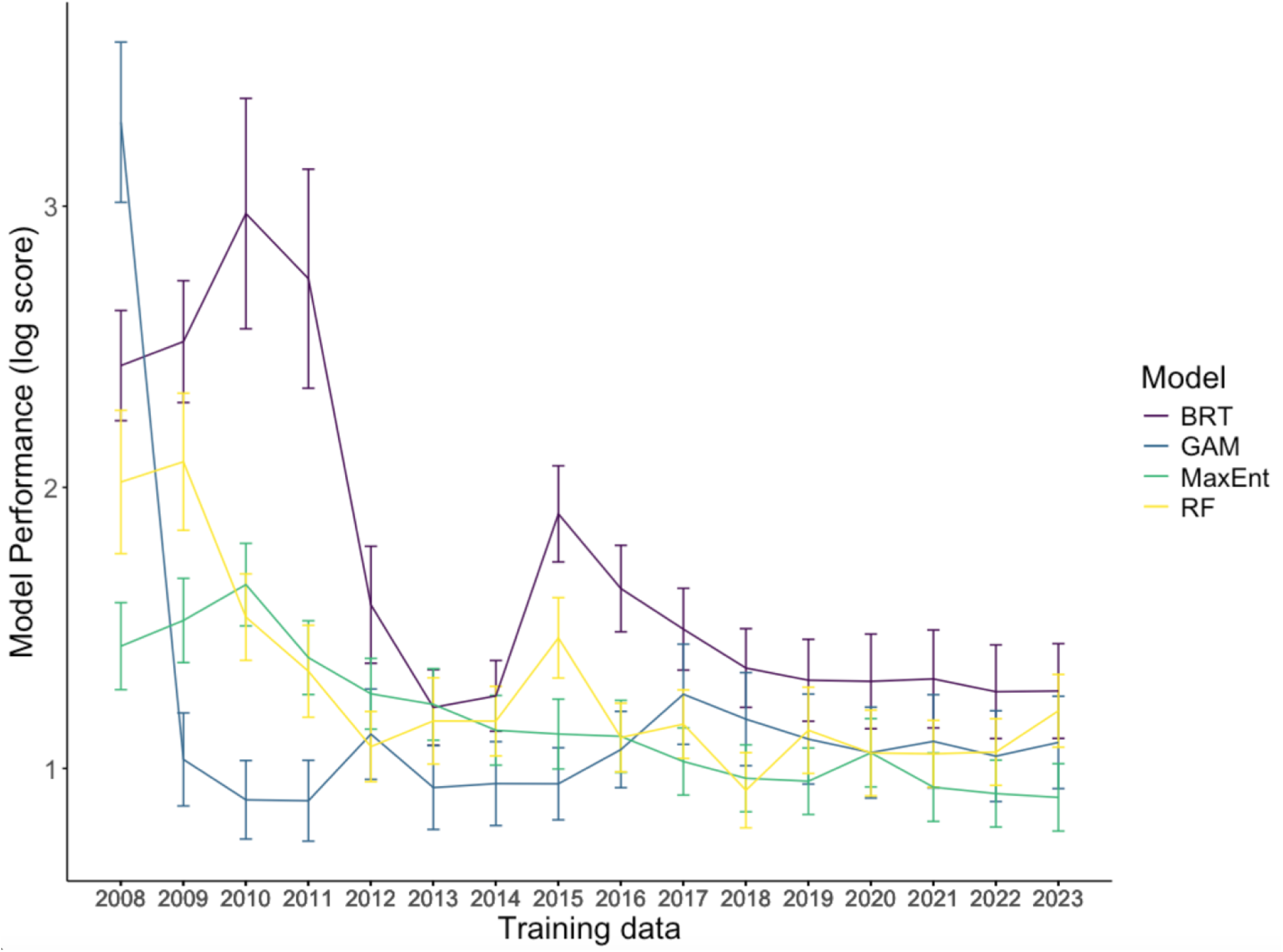
Performance of SDMs forecasting 2024 *Ae. albopictus* range with different amounts of historical training data. Effect of additional training data on the ability of each model (BRT = Boosted Regression Tree, navy; GAM = Generalized Additive Model, blue; MaxEnt = Maximum Entropy, green; and RF = Random Forest, yellow) to predict habitat suitability for *Aedes albopictus* (presence) in Suffolk County, New York, U.S.A. Year on the x-axis represents the most recent year of sampling added to the training dataset (i.e. 2012 indicates that all models were trained on data from 2008-2012, 2015 indicates models were trained on data from 2008-2015, etc.). Lines represent average log scores, and error bars represent ± 1 SE.

## Discussion

In this study, we used a 16-year (2008-2023) mosquito surveillance data from Suffolk County, NY, to evaluate a variety of SDM methods in predicting *Ae. albopictus* distributions across a heterogenous, seasonally shifting landscape. We identify key environmental drivers of distributions of *Ae. albopictus* and estimate habitat suitability spatially (across Suffolk County) and temporally (across mosquito season – May to October). We observed that habitat suitability in Suffolk County varied across space and across a mosquito season peaking in late summer, driven by environmental covariates such as impervious surface, greenness (EVI), temperature, and land use. Our analysis revealed that estimates of model predictive ability varied depending on the validation method used with MaxEnt and GAM performing better with field validation. Finally, we also observed GAM and MaxEnt had the highest predictive ability when using data from early in the invasion.

We found that overall habitat suitability in Suffolk County, New York was greatest in more urban areas (>20% Impervious Surface) for all models, while our models were split on the effect of vegetation – greenness (via EVI) – on suitability of habitat. Assigning different relationships between greenness and occurrence could be due to multicollinearity among predictors making it harder for models to define unique contributions of predictors. Many studies have found *Ae. albopictus* to be present in low to intermediate areas of impervious surface with vegetation present [34, 39, 54–56]. This aligns with what is known about *Ae. albopictus* as an ecologically flexible species that tends to dwell in suburban habitats where access to food sources and oviposition sites is high [57]. Male and female adult *Ae. albopictus* will feed on nectar which requires the presence of vegetation [58]. Female mosquitoes will feed primarily on mammalian hosts (humans, cats, dogs, etc.) [59] and oviposit in natural and artificial containers that tend to be in high availability in peri-urban spaces where there are natural sites (tree holes, etc.) and artificial (trash, lawn equipment, planters, etc.) [56].

Habitat suitability for *Ae. albopictus* in Suffolk County, NY, increased during the late summer. This is likely driven by temperature, an important climatic variable that determines fitness, precipitation, and relative humidity, although the latter two variables were not explored in this paper but have been shown to be important [37, 39, 49, 60, 61]. All models but one (BRT which assigned no significance), assigned positive relationships between daytime and nighttime land surface temperature and habitat suitability. This aligns with current understanding of the effects of temperature on the fitness of *Ae. albopictus* [49, 62]. Performance of many traits relevant to disease transmission and fitness have a unimodal relationship with temperature [62], increasing to an optimal temperature before declining rapidly to zero; however, these maximum temperatures, where fitness approaches zero, are not typically seen in Long Island, NY, which has a temperate climate [12, 49]. Under future climate warming scenarios, Suffolk County may see temperatures that are more unsuitable for *Ae. albopictus*, especially in more urban areas that are typically warmer.

Estimates of average habitat suitability in Suffolk County, New York, over a mosquito season (May - October) varied across SDM methods. Disagreement between modeling methods for predicting *Ae. albopictus* occurrence was the greatest in natural (forested, herbaceous, water, wetlands, shrub, barren) and altered (open – developed, cultivated) land classes. This is likely due to sampling bias in the training data. Ongoing vector surveillance in the area does not employ random sampling of sites; instead, because their goal is arthropod-borne disease surveillance, sites are chosen based on where they expect to see mosquitoes of interest (i.e. vectors of West Nile virus and eastern equine encephalitis) and where vector-borne disease cases are reported. As such, urban areas tend to be oversampled, and natural areas undersampled. This sampling bias likely contributes to the frequency of mispredictions among our SDMs [63]. SDM studies would benefit from data that are less spatially or environmentally biased; however, this is often not feasible. Instead, sampling bias correction methods can be leveraged to manage datasets that were collected with different goals [64].

In areas of disagreement, MaxEnt and GAM were most consistently accurate based on logarithmic scores across the field sampling period (July-September), despite GAM having lower out-of-sample AUC compared to the machine learning methods. The discriminatory ability of all models decreased in out-of-region (i.e. township sampling) cross-validation, suggesting they may have difficulty extrapolating occurrence in novel scenarios; however, this is not an uncommon issue as extrapolation can involve the introduction of new values for environmental covariates that models have not seen in training data [65, 66]. Interestingly, BRT might be assigning overly simplistic relationships between environmental covariates and habitat suitability as it assigned no significance to EVI, Daytime/nighttime land surface temperature, and land cover. Based on testing and training AUCs from the random sampling cross-validation, RF might have been overfitting as the model performs well on the training data, but quite poorly on the testing data. Improved predictive performance of RF and BRT in late mosquito season, as abundances of *Ae. albopictus* declined, was likely because both models tended to underpredict occurrence. GAM overpredicted occurrence throughout the season and MaxEnt both over and underpredicted. From a public health perspective, using model-informed surveillance will help target limited resources to areas predicted to be suitable for a given species. Further, using a model that overpredicts mosquito occurrence (GAM approaches) would better ensure sampling locations include the species of interest from a pest or disease control perspective. Collaborators at the Suffolk County Arthropod-Borne Disease Laboratory preferred BRT maps when presented with anonymized maps from all four models, as BRT had fewer areas of intermediate suitability, which made it easier to discern where to place mosquito traps. BRT maps also aligned well with their knowledge about the niche of *Ae. albopictus* in Suffolk County, New York.

A common difficulty with statistical or machine learning SDM approaches is predicting novel scenarios from current relationships between species occurrence and variation in environmental variables. This is especially important for using these approaches to predict future distributions of actively invading species like *Ae. albopictus*. One would expect that with the addition of more historical training data, models would continue to improve their performance in predicting near-term occurrence. When SDMs were trained using a modified expanding window approach and made to forecast occurrence into 2024 at sites of disagreement, model performance varied in response to the addition of new historic data across our SDM approaches. GAM and MaxEnt were the best at predicting future occurrence with the least amount of training data and remaining accurate as more years of data were added. While other models generally improved with the addition of more years of data, the addition of certain (e.g., 2015) years of data reduced predictive ability. The 2015 mosquito season may have had anomalous occurrence data due to extreme weather events, including a blizzard in January, which may have killed off diapausing eggs, and severe storms in August, that knocked out power for 63,000+ and likely affected mosquito abundance. GAM and MaxEnt performed the best when using data from early in the invasion to predict late-stage invasion distributions of *Ae. albopictus*. One study also showed that models trained on early stage invasion data could be used to predict later stage invasion data of an invasive wasp species in Europe [30]. Another study also found SDMs differed in predictive ability when predicting range expansion of deer species in Great Britain; however, these predictions were not better than a simple dispersal model [67].

While our study builds on growing literature exploring the applicability of SDMs to invasive species, there are several limitations. First, although our environmental covariates were chosen based on hypotheses we had about the niche space *Ae. albopictus* occupies, we did not include more host-specific variables (e.g., human population density), shown to be important in other studies [68–70]. Furthermore, some of our environmental variables (i.e. temperature) may have benefitted from downscaling as they likely operate on different spatial scales. Secondly, because the performance of Species Distribution Models is context-dependent, we recommend applying multiple modeling methods and comparing model performance to identify areas of weakness as in this study. Finally, while our general results should hold across multiple systems, the particulars (e.g., variable importance) may not transfer to other invasive mosquito species.

Overall, the statistical and machine learning models that we applied in this study performed reasonably well in predicting the distributions of *Ae. albopictus* and agreed on major patterns of habitat suitability, spatially and temporally. Our models predicted *Ae. albopictus* occurrence to be higher in moderate to developed habitat with high impervious surface and some vegetation as well as between the months of July to September. Disagreement among models at certain locations in Long Island for *Ae. albopictus* occurrence underscores the importance of using multiple methods when estimating species distributions and validating predictions with independent data, especially for invasive species that violate assumptions of equilibrium. While our mechanistic model performed poorly – if improved by accounting for variation in carrying capacity, for example – it could serve to predict distributions in places with no historic occurrence data, useful for predicting future invasive species occurrence with climate and land use change. Possible next steps could include incorporating time lags for temperature data and using temperature data collected at finer spatial resolutions to improve our mechanistic model. If used appropriately, SDMs can serve as powerful tools for predicting distributions of disease and invasive species in a changing world.

## Acknowledgments

We would like to thank the many seasonal interns at the Suffolk County Arthropod-Borne Disease Laboratory for their work identifying mosquito specimens and for collecting surveillance data for the past two decades.

## Supporting Information Captions

**S1 Figure.** Map of Suffolk County, New York, U.S.A., depicting the invasion of *Aedes albopictus* from 2008 to 2023. Sites where at least one female *Ae. albopictus* had been caught are pictured as points. Point color is based on the year *Ae. albopictus* was first detected. Sites that have never had *Ae. albopictus* are not pictured. Suffolk County shape file from NYS GIS Civil Boundaries Program (https://gis.ny.gov/civil-boundaries).

**S2 Figure.** Modified expanding window training approach to explore how the addition of more training data improves each SDM’s ability to predict late-stage invasion distributions (testing data). Each training iteration included an additional year of training data chronologically.

**S3 Figure.** Graphs display variation in a) % impervious surface, b) land cover, c) EVI, d) day surface temperature, and e) night surface temperature in Suffolk County, NY. These values are based on remote-sensed data averaged across May - October 2023. Color represents different values for each environmental variable. Temperature is represented in °C. Suffolk County shape file from NYS GIS Civil Boundaries Program (https://gis.ny.gov/civil-boundaries).

**S4 Figure.** Graphs represent receiver-operator curves generated from random sampling cross-validation. Each graph represents a different model. Each colored line represents a different iteration (different 20% testing and 80% training datasets, randomly selected) of cross-validation with bold black lines representing averages for testing (solid) and training (dashed) data. Text bubbles on each graph summarize the average AUC values for all training dataset iterations and all testing dataset iterations. As AUC gets closer to one, the model is considered to perform better. An AUC of one would indicate a model that can perfectly identify true presences and true absences.

**S5 Figure.** Graph represents receiver-operator curves generated from random sampling cross-validation for the mechanistic SDM based on temperature-dependent population density. Each colored line represents a different iteration (different 20% testing and 80% training datasets, randomly selected) of cross-validation with bold black lines representing averages for testing (solid) and training (dashed) data. Text bubbles summarize the average AUC values for all training dataset iterations and all testing dataset iterations. As AUC gets closer to one, the model is considered to perform better. An AUC of one would indicate a model that can perfectly identify true presences and true absences. AUCs close to 0.5 indicate that the model is no better at predicting presence or absence than a coin toss.

**S6 Figure.** Graphs represent receiver-operator curves generated from township sampling cross-validation. Each graph represents a different model. Each colored line represents a different iteration (different 20% testing and 80% training datasets, randomly selected) of cross-validation with bold black lines representing averages for testing (solid) and training (dashed) data. Text bubbles on each graph summarize the average AUC values for all training dataset iterations and all testing dataset iterations. As AUC gets closer to one, the model is considered to perform better. An AUC of one would indicate a model that can perfectly identify true presences and true absences.

**S7 Figure.** Partial dependence plots for seven covariates of interest: a) impervious surface, b) EVI, c) daytime land surface temperature, d) nighttime land surface temperature, e) Month, f) Year, and g) Land cover type that predict habitat suitability for *Aedes albopictus* in Suffulk County, New York, U.S.A. Points are colored by modeling method (BRT = Boosted Regression Tree, navy; GAM = Generalized Additive Model, blue; MaxEnt = Maximum Entropy, green; and RF = Random Forest, yellow). Suitability on the y-axis represents habitat suitability for *Aedes albopictus*.

**S8 Figure.** Graph represents total disagreement among our four species distribution models (Boosted Regression Tree, Generalized Additive Models, Maximum Entropy, and Random Forest) in predicting *Aedes albopictus* presence from July – September 2024 across different land use categories of Suffolk County, New York, U.S.A. Land use categories were determined from NLCD categories: natural (forested, herbaceous, water, wetlands, shrub, barren), altered (open – developed, cultivated), Developed – Low, Developed – Med/High (Developed – Med, Developed High). Higher values indicate higher disagreement among model predictions. Bars represent average disagreement. Letters indicate significant differences (α=0.05) calculated from an ANOVA followed by a Tukey post-hoc analysis.

